# Sparse neural networks enable low-power, implantable neural interfaces

**DOI:** 10.1101/2024.12.17.628936

**Authors:** Joseph T. Costello, Hisham Temmar, Luis Cubillos, Matthew J. Mender, Dylan M. Wallace, Madison Kelberman, Omer Benharush, Jeremy Simons, Matthew S. Willsey, Nishant Ganesh Kumar, Theodore A. Kung, Parag G. Patil, Cynthia A. Chestek

## Abstract

Recent advances in brain-machine interfaces (BMIs) using neural network decoders and increased channel count have improved the restoration of speech and motor function, but at the cost of higher power consumption. For wireless, implantable BMIs to be clinically viable, power consumption must be limited to prevent thermal tissue damage and enable long use without frequent charging. Here, we show how neural network “pruning” creates sparse decoders that require fewer computations and active channels for reduced power consumption. Across multiple movement decoding tasks using brain and muscle signals, recurrent neural network decoders can be compressed by over 100x while maintaining strong performance, enabling decoding on the implant with <1% power increase compared to decoding externally. Pruning also allows for deactivating up to 89% of channels, reducing BMI power by up to 5x. Counterintuitively, our findings suggest that BMIs employing a subset of a larger number of channels may achieve lower power consumption than BMIs with fewer channels, for a given performance level. These results suggest a path toward power-efficient, implantable BMIs suitable for long-term clinical use.

## Introduction

Neural interfaces show promise in restoring mobility and communication to people with sensorimotor impairments. Brain-machine interfaces (BMIs), particularly intracortical BMIs using electrodes implanted in the cortex, have enabled accurate control of robotic arms^1^ and virtual quadcopters^2^, re-animation of paralyzed limbs through functional electrical stimulation^3^, high speed cursor control^4^, and speech decoding approaching able-bodied levels^5,6^. Outside the brain, implanted electromyography (EMG) and peripheral nerve interfaces have allowed amputees to control prosthetics with increasing dexterity and stability over time^7–9^. Recently, the use of artificial neural networks for decoding neural activity has led to significantly higher performance of real-time BMIs and EMG interfaces^2,5,6,10^. Additionally, there are many efforts to develop BMIs that can record from thousands of implanted electrodes^11–13^, which may further improve decoding accuracy.

However, with goals of developing widely accessible, implantable BMIs, power consumption may be a major limiting factor as BMIs expand to more complex algorithms and hardware.

Implantable wireless interfaces are typically powered by an internal battery and use wireless power transmission for recharging the battery. The total power available to the system is thus limited by the battery capacity and safe thermal radiation limits of the body. For devices near the brain, tissue must not be heated more than 1-2°C to remain safe^14^, limiting on-device processing and the strength of wireless power. For the widespread adoption of wireless battery-powered BMIs, the system would ideally have a small battery and last for at least one day without recharging, ideally longer.

To process neural signals, BMIs use power for amplifying voltage recordings, filtering, sampling, and feature extraction (Figure 1). Of these, the amplifier may consume over 90% of the total system power^15^, where power consumption scales approximately with the number of recording channels. Recent developments in integrated chip design have resulted in increasingly power efficient chips for neural recording that use <3uW per channel^16–19^, and neural features like spiking band power can reduce power usage by lowering the sampling rate by 5-10x^20^. An alternative method is to simply turn off non-informative recording channels to remove their associated power consumption. Previous methods have shown that 57-88% of channels can be deactivated for less than 5% loss in decoding accuracy^21–26^, but have not been evaluated on high channel count systems. An ideal channel selection method would easily incorporate into the existing decoder training and work with any decoder architecture. Importantly, the ability to deactivate many channels does not imply that fewer electrodes can be implanted, as the initial quantity is necessary to identify the most informative channels. The benefits of having more electrodes and being able to down-select channels for extended battery life should be weighed against the potentially increased surgical risk with implanting more electrodes^27^.

**Figure 1:**
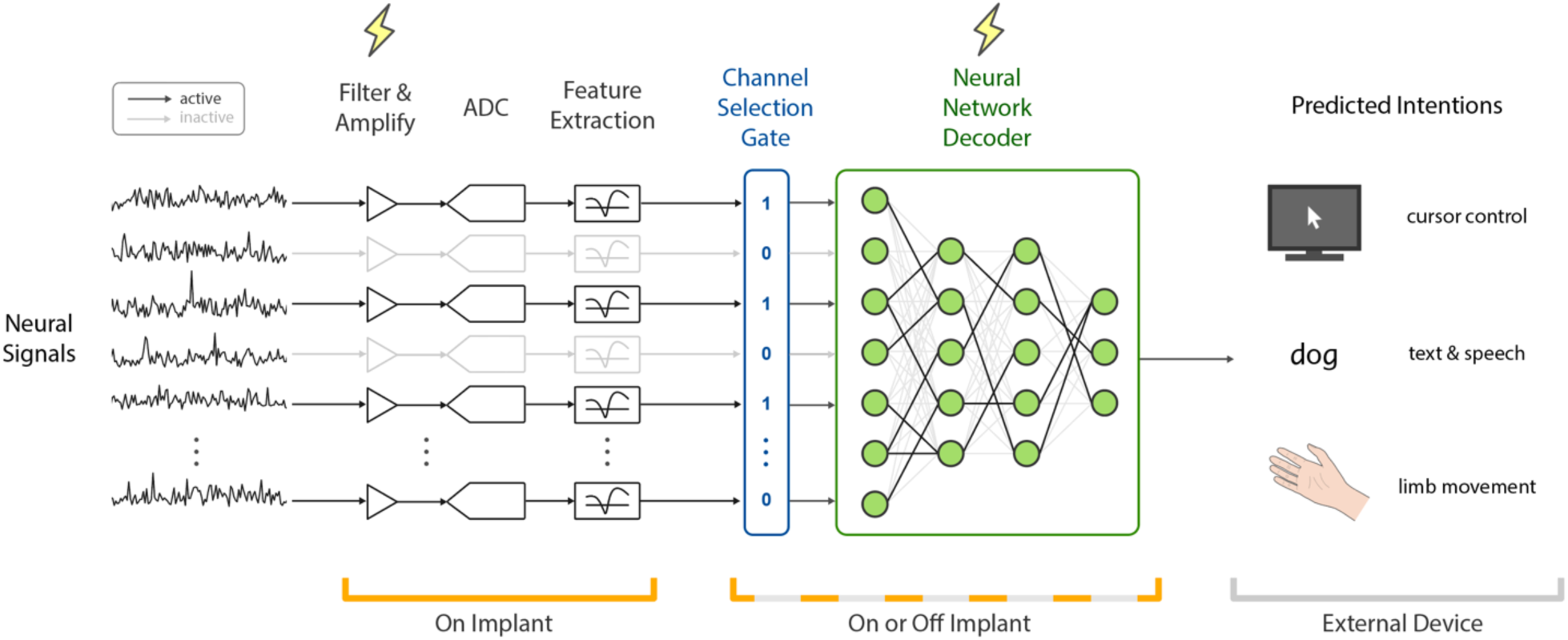
An implantable neural interface using a sparse neural network decoder. Neural voltage signals are recorded from electrodes by performing filtering and amplification, conversion to digital samples (“ADC”, analog-to-digital converter), extracting features for decoding, and decoding with a neural network to predict the user’s intentions. The amplifier and decoder have significant power consumption. We introduce a sparse channel selection gate (blue) that allows for many of the input amplifiers to be turned off to save power. The neural network decoder (green) can be compressed using pruning to have relatively few parameters and low power consumption, providing the option of running directly on the implant instead of on an external computer.

After neural features are extracted, a decoding algorithm is used to predict user intentions (Figure 1), which can be performed on or off the implanted device. Due to significant computation and power requirements, BMIs with neural network decoders often rely on GPUs outside the body^5,16,28,29^. On the implanted device, power is required for wirelessly transmitting neural features in real-time, using data rates as high as 200 Mbps (sending all samples for 1000 channels, 20 kHz, 10 bit resolution). However, if the decoder can be significantly compressed with minimal memory usage and number of computations, it could be run directly on the implant for reduced wireless bandwidth and lower latency. Previous work has shown how network “pruning” can compress networks by >10x without a significant drop in accuracy^30–32^. With pruning, small magnitude network weights are set to zero, removing their associated multiplicative operations and storage in memory, resulting in a sparse network. However, it is unclear how well these compression methods may perform for BMI decoders, and their robustness to distribution shifts during closed-loop control^33^.

Here we show how neural network pruning can reduce the overall power consumption of implantable neural interfaces. Using pruning, we find that high performing recurrent neural network decoders can be compressed by over 100x, for both EMG and intracortical decoding, for a similar number of parameters to linear decoders. In closed-loop decoding experiments with non-human primates, sparse decoders had no significant performance drop and can be run on low-cost embedded devices. Then, by adding a simple channel selection layer to neural network decoders, we show how pruning can select the most important neural channels across a variety of decoding tasks and number of active channels. Counterintuitively, BMIs with more channels may be able to achieve lower power consumption than systems with fewer channels by selecting a smaller subset of more informative channels. By adding multiple channel selection layers, decoders can use varied numbers of channels, enabling the option for a “battery saving mode” for BMI. These methods aim to reduce power consumption constraints, allowing for greater flexibility in the design of neural interfaces.

## Results

### Reducing neural network power consumption using pruning

With the goal of reducing the number of operations performed by the decoder, we investigated the ability of pruning to compress neural network decoders of varied architectures. To prune the decoder network, we iteratively zero-out the smallest magnitude weights during decoder training^32^ (Figure 2a), which removes their associated multiplication and addition operations. To prune a slightly greater fraction of weights without performance loss we apply “pruning with rewinding”^34^, in which the final set of sparse weights are reset to their initial values and retrained multiple times (Figure 2a; see Methods). As an initial example, a long short-term memory (LSTM) decoder of finger movement could have up to 98% of weights pruned for less than 1% increase in mean-squared error (MSE) for velocity and position predictions, requiring only 5.6k out of the original 364k parameters (Figure 2b). To account for the removal of pruned weights, the remaining weights tend to have larger magnitudes (Figure 2b, bottom).

**Figure 2:**
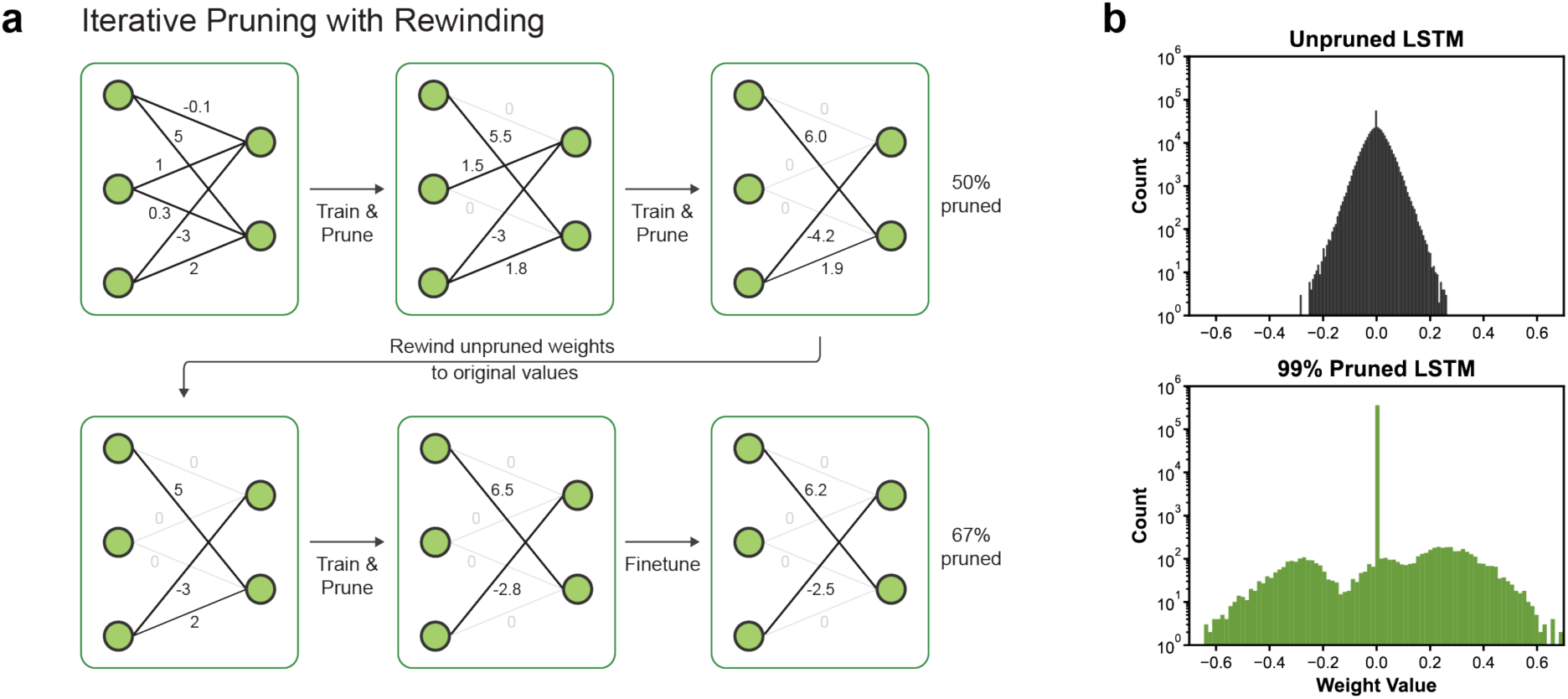
Neural network pruning reduces the number of decoder computations. (a) Toy example of pruning with rewinding. At each step, network weights are trained (updated) and then the lowest magnitude weights are pruned (set to zero). After reaching a chosen pruning amount (here 50%), the unpruned weights are reset back to their original values before another round of training and pruning. This “rewinding” can be repeated multiple times. The first pass of training/pruning allows the network to learn which weights are important, and the proceeding rewinding/training/pruning allow the model to fine tune those specific weights for improved performance. (b) Histogram of parameter values for an unpruned LSTM (top) and a pruned LSTM with 98% of weights set to zero. Parameters shown are after training has completed, for predicting 2-DoF finger movement from 96 intracortical channels. The pruned decoder has only 1% increase in MSE in this example.

First, we investigated the maximal degree of decoder compression across multiple movement decoding datasets and generic decoder architectures. We tested the LSTM (mentioned above), a simple recurrent neural network (“RNN”), and a temporal-convolutional feedforward network (“TCN”), of which the LSTM and TCN have previously demonstrated strong real-time decoding performance^28,35^. To establish a baseline, we trained minimally sized networks optimized for offline accuracy, with the LSTM and RNN limited to a single layer, and containing an order of magnitude fewer parameters than recent offline models^36–38^. Then, we found the minimum number of parameters needed for a given performance level by sweeping over varied pruning percentage (0 to 99%) and network size. We performed this analysis for decoding movement with the following datasets (see Methods for more details):

- 2-DoF finger movement from 96-channel intracortical signals in an NHP on 10 days (Figure 3a/e)
- 2-DoF finger movement from 8-channel implanted EMG in an NHP on 5 days (Figure 3b/f)
- 2-DoF arm reaches from 130-channel intracortical signals in an NHP on 1 day (Figure 3c/g)
- 5-DoF movement from simulated 1000-channel intracortical signals for 5 simulations (Figure 3d/h)

**Figure 3:**
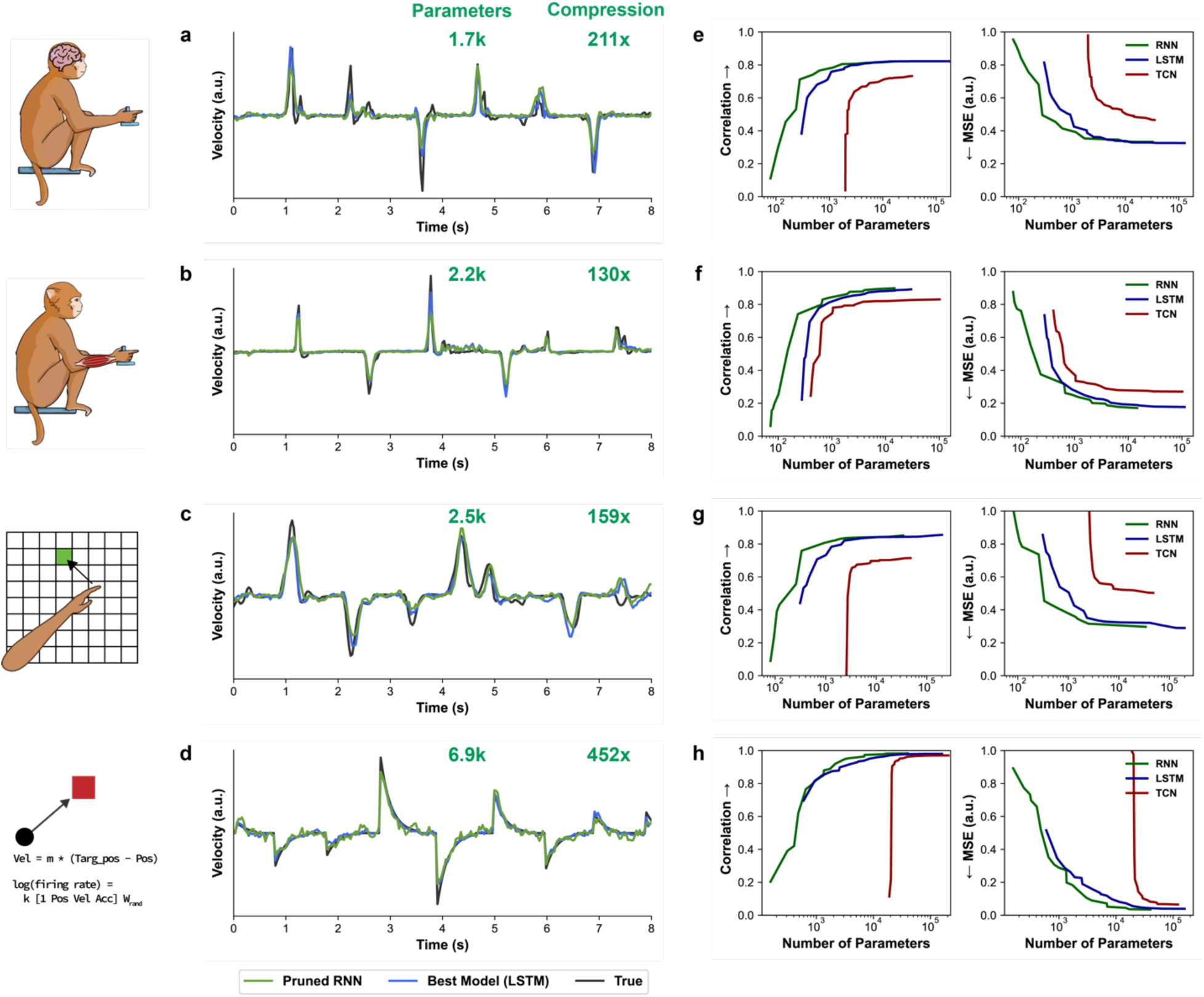
Pruning significantly compresses neural network movement decoders. (a-d) Example velocity predictions for continuously decoding (a) finger movement from 96-channel intracortical signals, (b) finger movement from 8-channel EMG, (c) arm reaching from 130-channel intracortical signals, and (d) simulated 5- DoF movement from 1000-channels. The true movement is shown in black, the best baseline (unpruned LSTM) model predictions in blue, and the pruned RNN predictions in green. The pruned RNN has less than 0.02 decrease in correlation compared to the baseline model. (e-h) Minimum kinematic prediction error for a range of model sizes and 3 architectures. For each dataset and model architecture, the model size and pruning level was varied, and accuracy averaged across datasets. The lines shown represent the highest correlation or lowest MSE for a given number of parameters (a Pareto boundary).

Pruning each decoder type allowed for a multiple order of magnitude reduction in the number of parameters (Figure 3). For example, on the intracortical-finger dataset, the default LSTM achieved the highest average kinematic correlation (0.82) of all three architectures using 364k parameters, which we used as a baseline parameter count. The best pruned LSTM used 2.3k parameters for 141x compression (95% pruning with 4x smaller hidden size), for less than 0.02 decrease in average correlation. Notably, the best pruned RNN required only 1.7k parameters for the same accuracy level, for 211x compression (95% pruning with 128 hidden size). For comparison, a linear Kalman filter (as in ^4,39^) used 985 parameters (96 channels using steady-state gain).

The number of parameters and compression ratio for less than 0.02 decrease in correlation relative to the baseline LSTM for each dataset are shown in Figure 3a-d. Pruned RNNs enabled the greatest compression ratios of the network architectures, ranging from 130-452x across datasets. As shown in Figure 3e-h, further reductions in parameter count for any architecture can be achieved if slightly lower accuracy levels can be tolerated.

Additionally, we observed that pruning 50-90% of weights resulted in an improvement in MSE compared to unpruned models for the RNN (Supplemental Figure 2). This suggests that pruning acts like a beneficial regularizer, enabling the RNN to achieve comparable MSE to the LSTM with fewer parameters. With fewer computations, this makes the pruned RNN a strong candidate for running on a low-power embedded device.

### Decoding finger movement in real-time using sparse decoders

Next, we evaluated the performance of sparse (pruned) decoders for closed-loop control. With significantly fewer parameters, it was unclear if sparse decoders would be better regularized or if they would be less robust to shifts in the input distribution that may occur during closed-loop control^33^. On 7 closed-loop comparison days, Monkey N used both a sparse and non-sparse decoder (either LSTM or TCN) to perform a continuous finger target acquisition task, using either 96-channel intracortical signals or 8-channel implanted EMG as the neural input.

Grouping all trials across 7 test days, the sparse decoders were 3.7% faster on average with no significant difference compared to each day’s non-sparse decoder (p=0.04, two-tailed t-test with Bonferroni correction). Within these test days, the sparse decoders had similar offline MSE compared to non-sparse decoders, ranging from 11% lower to 32% higher compared to the non-sparse decoders (Figure 4a; note that sparse decoders had varied degrees of pruning). Online, the sparse decoders had similar trial times, ranging from 16% faster to 14% slower, where the sparse decoder was significantly faster on only one day and significantly slower on only one day (Figure 4b; p<0.005, two-tailed t-test with Bonferroni correction). For 6 out of 7 days, lower offline MSE resulted in faster online trial times. These results demonstrate that the sparse decoders had no significant loss in real time performance.

**Figure 4:**
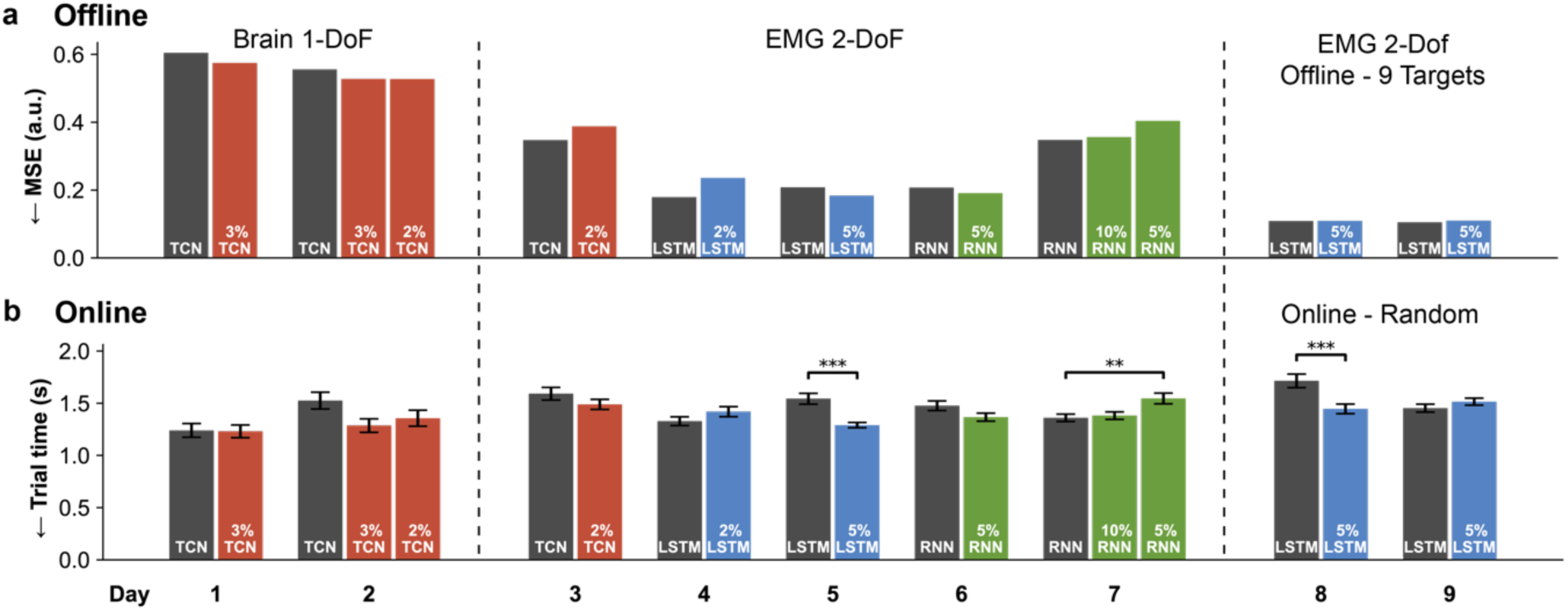
Closed-loop finger movement decoding with pruned decoders. (a, b) Offline (open-loop) and online (closed-loop) performance of non-sparse (grey) and sparse (green) decoders, where each group of bars is a separate experimental day and the type of signal & task is specified at the bottom. A “2% LSTM” means that 2% of parameters are non-zero, and 98% are pruned to zero. Data are from Monkey N performing a random- target finger task. In the right-most two sessions, the decoder was trained on only 9-targets but tested online with random targets, as a test of generalization. (a) Offline MSE between true movement and predicted movement, averaged across position and velocity predictions and across DoFs. (b) Online trial times, averaged across the 150-400 trials of each decoder. Significance between decoder trial times was calculated using a two-tailed t-test with Bonferroni correction and are indicated as follows: **p < 0.01, and ***p < 0.001. Error bars show 1 S.E.M.

To test generalization to new movements, we trained both a non-sparse and a sparse LSTM on a 9- target task and then tested each decoder online on a random target task with the full range of target locations (EMG decoding). Both the non-sparse and sparse LSTM had similar MSE for the 9-target training dataset (<0.01 a.u. difference). When tested in the full task online, compared to the non-sparse decoder, the sparse decoder had significantly faster trial times on one day (1.4 vs 1.7 sec, p<0.005, two-tailed t-test) and slightly, but not significantly, slower but trial times on a second day (1.5 vs 1.4 sec respectively, p=0.23, two-tailed t-test). Thus, the sparse decoder had no significant loss in generalization ability.

To run neural decoders on an implant without the need for customized hardware, we developed an embedded-C code library for efficiently running sparse networks trained in PyTorch. Using a low-cost Arm-M7 microcontroller, a pruned RNN with 1.7k parameters decoded 96-channel intracortical data with a latency of 1.72 ± 0.01 ms. This computation time is well within the 32 ms bin-size and comparable to decoding with a non-compressed network on a GPU^28^.

### Reducing power consumption through sparse channel selection

We next investigated the ability of pruning to perform channel selection, with the goal of maintaining strong decoding accuracy with few active channels (for lower power consumption). To do so, we added an input gate layer to the start of an LSTM decoder that pruned (deactivated) the least important channels. We used the previously mentioned datasets with intracortical recordings and also tested pruning channel selection for decoding human speech (dataset from Willett et al. 2023). For each dataset we measured the maximum number of channels that could be removed for <5% drop in correlation compared to using all channels, or <5% increase in phoneme error rate for the speech dataset.

Decoder velocity predictions using few active channels qualitatively matched true velocities across tasks (Figure 5a-c). As in Figure 5e-h, large reductions in the number of active channels could be achieved: 2-DoF finger movements – 89% reduction, 2-DoF reaching – 81%, simulated 5-DoF 1000 channels – 89%, speech decoding – 50%.

**Figure 5:**
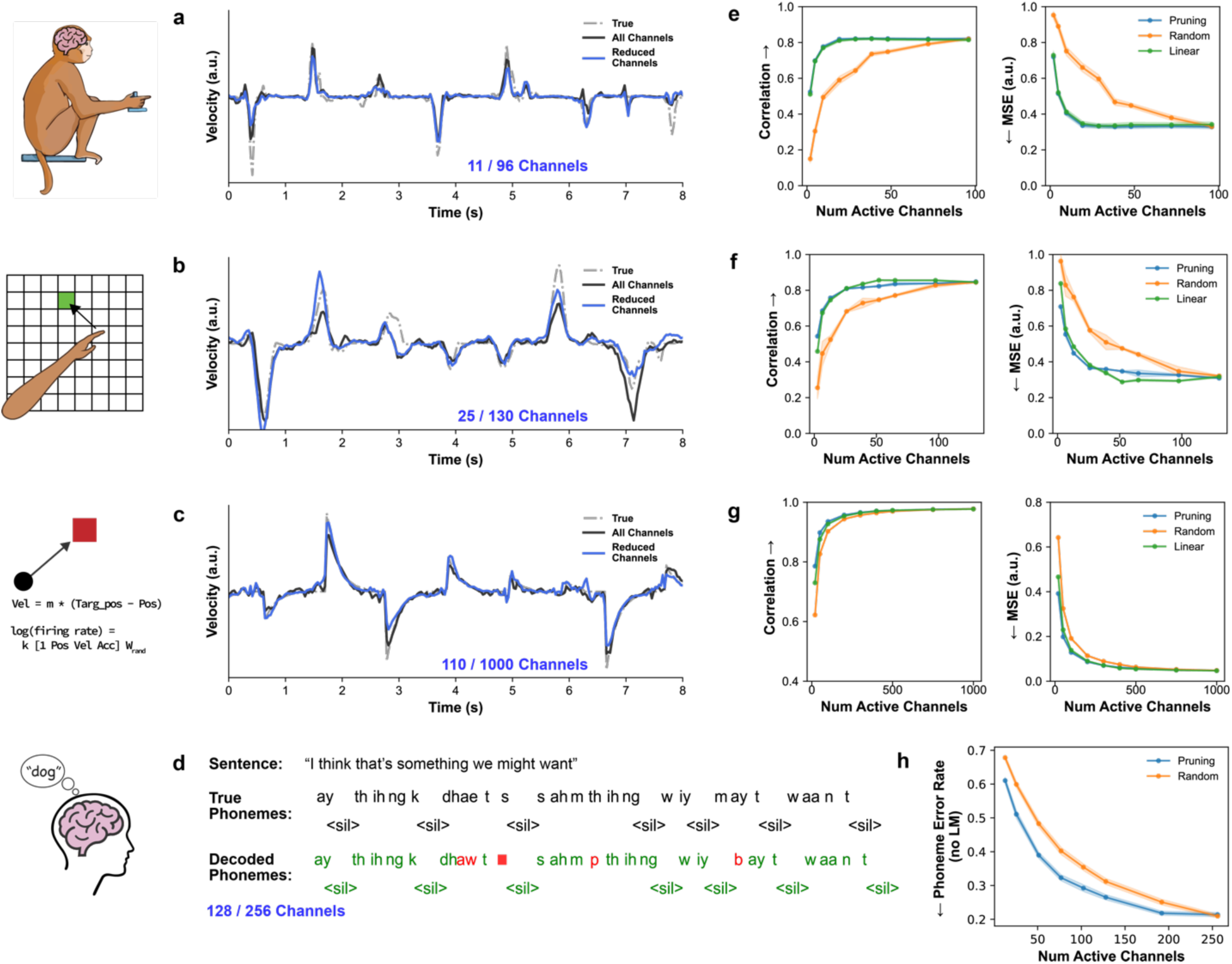
Channel selection using pruning. (a-c) Example velocity predictions using few active channels for decoding (a) finger movement from 96-channel intracortical signals, (b) arm reaching from 130-channel intracortical signals, (c) simulated 5-DoF movement from 1000-channels, where the chosen number of channels corresponds to a less than 5% decrease in compared to using all channels. (d) Example decoded speech phrase using few active channels from 256-channel intracortical signals, where the chosen number of channels corresponds to a less than 5% increase phoneme error rate compared to using all channels. Correctly decoded phonemes are in green, incorrect in red, and ‘ <sil>’ indicates the silence token for the end of a word. Phonemes were classified by taking the most likely phoneme at each time step (no correction by a language model). (e-h) Decoder accuracy vs number of active channels, for pruning-based selection (blue), linear correlation selection (green), and random channel selection (orange). In ‘linear’ selection, the channels with the highest correlation with velocity were selected. Error bars represent 1 S.E.M. across datasets.

As a baseline, we compared pruning-based selection to random and linear channel selection. For the three movement decoding tasks, at the previously mentioned fraction of active channels, pruning-based selection had significantly higher correlations compared to using random subsets of channels (p < 0.001, two-tailed t-test). Compared to simple linear selection (see Methods), pruning had significantly higher correlation for the simulated dataset (p < 0.001, two-tailed t-test) and no significant difference (p > 0.05, two-tailed t-test) for the others.

### Lower BMI power consumption using higher channel-count arrays

With higher channel count BMIs, there is higher total power consumption compared to lower channel count systems when all channels are active. However, in a higher channel count system with a larger pool of channels to pick from, there are likely more strongly tuned, informative channels than a lower channel count system with a smaller starting pool (assuming the same distribution of channel tuning). We hypothesized that for a given accuracy level, higher channel count BMIs could use fewer active channels to reach the accuracy level using lower power consumption. Figure 6a provides a toy example, in which we consider each channel to have low, moderate, or high information with a consistent distribution between a small array and a large array. The large array has more high-information (green) channels simply because it has more channels overall and can reach the desired (arbitrary) decoder accuracy using just those channels. However, the small array requires the use of many moderate-information channels to reach the desired accuracy, using more channels overall.

**Figure 6:**
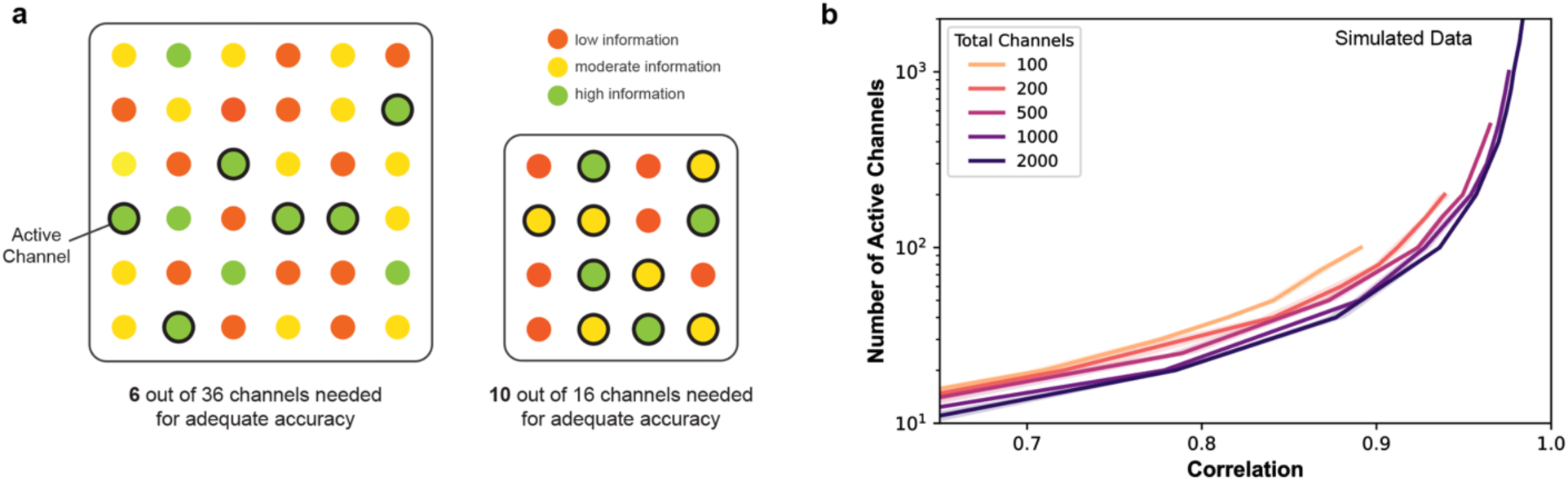
Lower power consumption when using larger arrays. (a) Toy example of selecting channels in a larger and smaller array (arbitrary data). We assume there is some distribution of channels that contain low/moderate/high information for the desired decoding task, and that we have a desired level of decoding accuracy. With the larger array, only 6 channels are needed to achieve the desired accuracy since there are more high information (green) channels. With the smaller array, there are fewer green channels so more moderate-information (yellow) channels are needed to hit the desired accuracy, using more active channels and more power overall. (b) Number of active channels needed for a desired accuracy (correlation), using simulated datasets. With fewer total channels (lighter traces), more active channels and more power is needed. Correlation is between predicted and true kinematics for a 5-DoF task. Error bars denote 1 S.E.M, averaged over 3 simulation runs.

We tested this concept using simulated datasets with the total number of channels ranging from 100 to 2000, each with random tuning, and trained gated LSTM decoders with varied numbers of active channels. As in Figure 6b, we found the number of active channels required for a given accuracy level, as measured by the correlation between predicted and true kinematics (5-DoF task). As expected, for a constant total number of channels, more active channels were needed to reach higher accuracy levels. However, for a given accuracy level, fewer active channels are required when the total number of channels is higher, as seen by the darker traces below the lighter traces in Figure 6b. For example, to achieve a 0.85 correlation, a 100-channel system needs 58 active channels on average while a 2000-channel system only needs 34 active channels on average, representing a 41% reduction in active channels. In this simulation, however, there may be a ceiling to this improvement, as power savings between a 1000 and 2000 channel system were much smaller (39 vs 34 channels required for 0.85 correlation, respectively). While the data here is simulated and may not match the real number of channels needed for a given correlation, it suggests that with more starting channels lower power consumption can be achieved by using fewer active channels.

### A “low power” mode for implantable BMI

Finally, we aimed to develop a single decoder that could work with varied numbers of active channels, allowing for varied power consumption. While one could train multiple (separate) decoders for each number of active channels (as was done in the previous sections), this may be costly in terms of the training time and the memory/storage required for a device to switch between decoders. We found that simply adding separate channel selection gates with an affine layer for each number for active channels, with a single decoder network, resulted in high decoding accuracy (Figure 7a). This multi-gate model had only 0.005 drop in correlation on average compared to separate decoders trained on each number of active channels (Figure 7b; simulated 1000-channel 5-DoF task). Thus, adding multiple channel selection layers enables a “low-power” mode where one can have significant reduction in power for a small reduction in accuracy.

**Figure 7:**
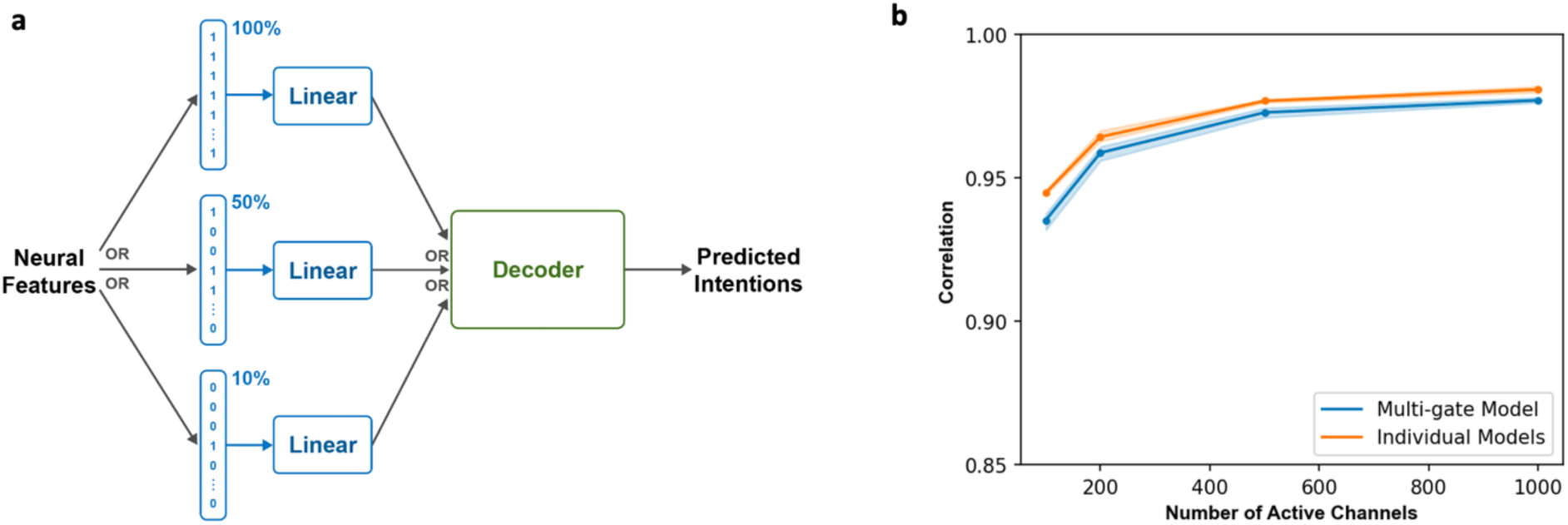
Modulating BCI power using a multi-gate decoder. (a) Architecture overview. Neural features are passed through the desired channel gate and linear layer, which determines which channels can be turned off to save power. A single “core” decoder works with any of the gates. (b) Performance of the multi-gate model compared to training individual decoders for each number of active channels. Decoders were trained on a simulated 1000-channel dataset and performance averaged over 3 repetitions. Error bars represent 1 S.E.M.

## Discussion

BMIs using artificial neural network decoders and higher channel counts have recently demonstrated performance approaching able-bodied control^6,28,35^, but at the cost of increased computational complexity and power consumption. For translation to high performance, fully implantable BMIs that function for long periods on a small battery, power consumption is a limiting factor. Here we showed how neural network pruning can reduce BMI power usage and has implications for the design of the entire BMI. We found that multi-DoF movement decoders could be compressed by over 100x, for a total parameter count under 7k, which is on the same order of magnitude as a linear decoder. Using pruning for channel, we found that 50-89% of channels could be deactivated (powered off) without significant performance loss, for approximately 1.8-5x power savings (see Supplement). Both results follow the “lottery ticket hypothesis”^40^, which suggests a benefit in finding an optimal subset within a larger pool rather than starting with a smaller pool.

Current BMIs using neural network decoders perform decoding on an external computer^5,6,16,28,36^. Alternatively, decoding could be performed on the implanted device if the decoder can be compressed for low power consumption. Here we demonstrated a high performance RNN decoder running in real-time on a low-cost microcontroller, where sparse decoders may be run using <0.3% of the full system power (see Supplement). Further reductions in computation (and power usage) could be achieved by quantizing parameter values to fewer bytes^41^.

Running the decoder on-device may allow for lower latency by significantly reducing the amount of data wirelessly transmitted from the implant (by transmitting control signals rather than neural data) and removes the need for a separate processor in the loop when controlling external devices. This also improves the privacy and security of neural data since data stays on the implant and is never stored on external computers or servers. However, on-device decoding may not make sense if the decoder needs frequent updating, or if neural data can be significantly compressed for low data-rate transmission (see Supplement for additional considerations). Future work may explore the ability of sparse decoders to generalize across tasks. Interestingly, we found that pruned decoders showed no loss in ability to generalize to additional movements in the 2-DoF finger task. Offline, RNN decoders with pruning had improved accuracy compared to the non-pruned decoder when 50-90% of weights were removed (Supplemental Figure 2). This may suggest pruning has a regularization effect similar to neuron dropout.

We demonstrated how neural network pruning can also perform channel selection, with the goal of maintaining decoding accuracy with a minimal set of channels. While pruning had no significant improvements over simple linear channel selection in 2 out of 3 datasets, it can easily be added to existing neural network training and works with any existing decoder loss function for regression or classification tasks. Channel selection allows for the tradeoff between decoding accuracy and power consumption, where BMIs could actively modulate the number of active channels to balance this tradeoff. If a lower decoder accuracy can be tolerated by the user, a large proportion of channels could be deactivated to significantly extend the battery life (like the power saving mode on mobile phones). The number and specific subset of active channels could also be varied based on task, where neural activations and important channels may vary with small task changes^42^. Higher complexity tasks (like controlling many degrees of freedom) may need more channels. In an additional offline analysis, we found that few-channel decoders maintain a similar noise robustness (Supplemental Figure 3). While dimensionality reduction techniques like principle component analysis can reduce the number of features needed for decoding, they still require all channel amplifiers to be active, for limited power savings. Future work may validate few-channel decoders for real-time control and determine if the decoder needs more frequent updating to account for channel activity drift or task changes^2,42–44^.

Using simulated data, we found that BMIs using a subset of a larger number of channels could use fewer active channels (and less power) than systems with fewer total channels. Intuitively, larger electrode arrays may be more likely to have more highly informative channels simply because of the larger starting pool, and one can use channel selection to activate the most informative channels. This observation lends to the strategy of implanting as many electrodes as safely possible so the BMI can achieve maximal accuracy or minimal power usage at reduced accuracy. Toward this goal, there are many recent efforts to develop high channel count arrays^11–13,45^.

However, this strategy should be carefully weighed against strategies of developing electrodes with improved neuron yield^46^ or improving brain targeting for better decoding^1,6^ which may not have the potentially increased surgical risk and complexity for implanting more electrodes^27^.

## Acknowledgements

We thank Eric Kennedy for animal and experimental support. We thank the University of Michigan Unit for Laboratory Animal Medicine for expert surgical and veterinary care. This work was supported by NSF GRFP 1841052, NSF EFRI BRAID 2223822, NSF NCS 1926576, NIH R01-NS-105132-01-A1, NIH U41 NS129436, and the Agencia Nacional de Investigacion y Desarrollo (ANID) of Chile.

## Author Contributions

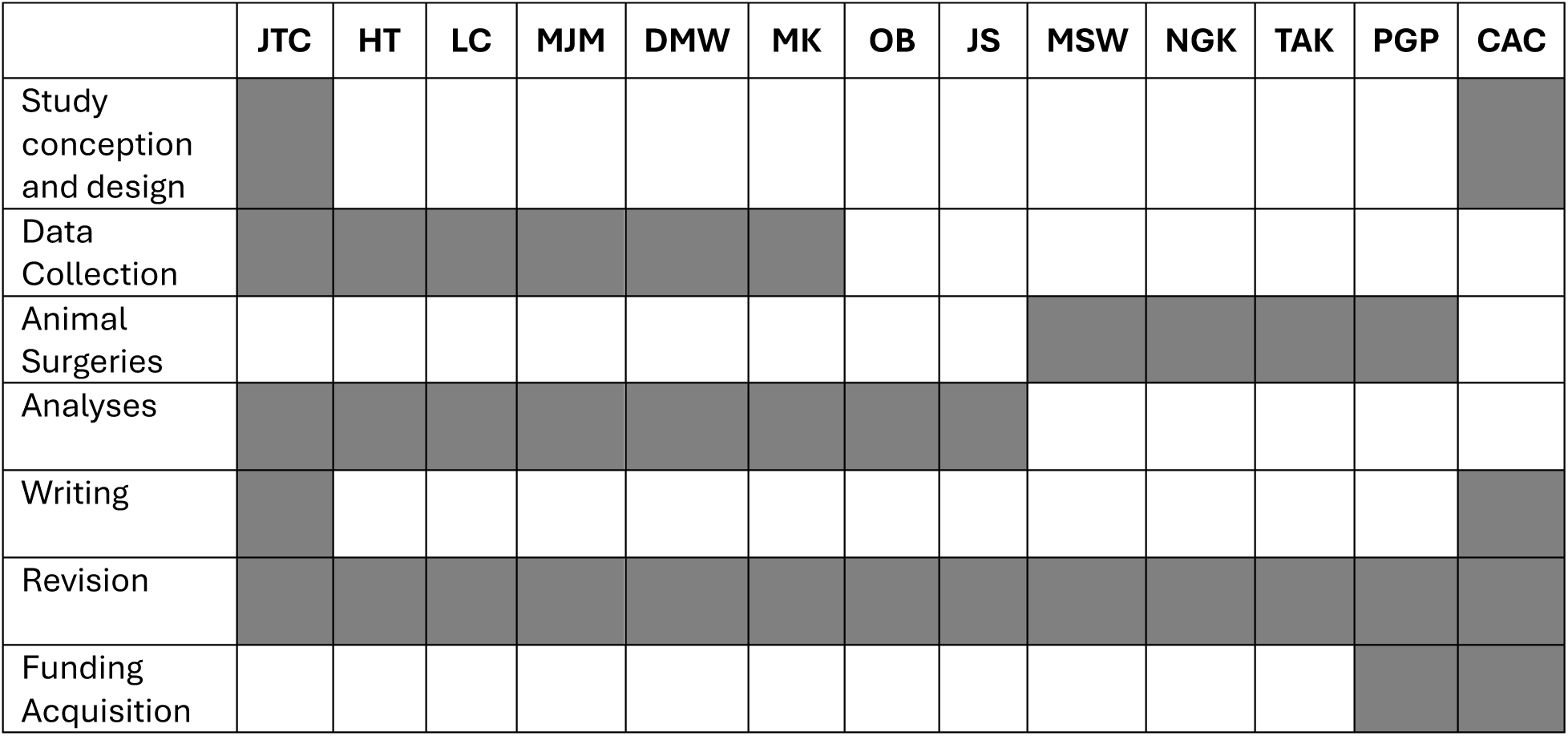

## Code Availability

Code is available at: github.com/jtcostello/sparsebmidecoders

## Methods

### Neural Datasets

We tested decoders on several neural decoding datasets with varied tasks, type of neural signal, and number of channels.

### Finger Movement Task (Brain and EMG recordings)

To test the ability of intracortical brain signals and EMG signals to decode fine finger movement, we implanted one male rhesus macaque (Monkey N) with multiple electrode types and trained him to perform a finger movement task. Monkey N was implanted with Utah microelectrode arrays (Blackrock Neurotech) in the primary motor cortex (M1) using the arcuate sulcus as an anatomic landmark for hand area, as described previously^47,48^. All 96 channels in M1 were used for closed-loop BMI control and channel selection analyses, and a subset of the channels with threshold crossings morphologically consistent with action potentials were used for offline pruned decoder analyses. In a subsequent surgery, Monkey N was also implanted with chronic bipolar intramuscular EMG recording electrodes (custom PermaLoc electrodes, Synapse Biomedical. Inc.) in the muscle belly of 8 muscles controlling the fingers and wrist as outlined previously^49^. The location of each lead was determined by intraoperative stimulation in conjunction with anatomical landmarks. All 16 EMG channels (8 bipolar pairs) were used for offline recordings and online closed-loop control. Surgical procedures were performed in compliance with NIH guidelines as well as our institution’s Institutional Animal Care & Use Committee and Unit for Laboratory Animal Medicine.

Neural signals, either brain or EMG on each day, were recorded using a Cerebus neural signal processor (Blackrock Neurotech). For brain recordings, the Cerebus sampled the 96 channels at 30 kHz, applied a 300-1000 Hz bandpass filter, and downsampled to 2 kHz to calculate spiking band power^20^. For EMG recordings, the Cerebus recorded the 16 channels of the 8 bipolar leads where each bipolar pair had one contact referenced to the other to reduce noise, sampled data at 30 kHz, applied a 100-500 Hz bandpass filter, and downsampled to 2 kHz. The resulting signal values (brain or EMG), were streamed to a running xPC Target version 2012b (Mathworks), which took the magnitude of the signal and summed into 32 ms bins.

We trained monkey N to acquire virtual targets with virtual fingers shown on a computer screen in front of the animal. In hand control (“offline” trials), the monkey moved his fingers within a manipulandum that measured the angles of the index and middle-ring-small (MRS) finger groups using bend sensors, which controlled the virtual fingers. During brain control (also referred to as closed-loop, “online” trials), the monkey controlled the virtual fingers using neural signals and a decoder. The finger task required placing the virtual index finger and MRS fingers on the respective targets and holding for 750 ms during offline trials or 500 ms during online testing. The target size was 15% of the active range of motion and randomly placed at the start of each trial. Some sessions used 1 target (1-DoF), in which the monkey had to move all fingers together, while others used 2 targets (2-DoF, index and MRS fingers independently).

We collected offline data with 500 trials across 10 brain recording sessions over six months and 5 EMG recording sessions over 3 months. We tested online brain decoding on 2 sessions and online EMG decoding on 5 sessions.

### Reaching Task

For a representative computer control task with naturalistic, self-paced movements, we used the MC_RTT dataset from the Neural Latents Benchmark^50^. Briefly, one rhesus macaque monkey performed sequential reaches to randomly placed targets on an 8x8 grid. Neural activity was recorded from a 96-channel electrode array in motor cortex (M1) and 130 sorted units were extracted. Here, sorted unit activity was binned into 32 ms bins and was used to predict x/y finger position and velocity. Data was from a single recording session with 649 seconds and 789 trials (reaches).

### Simulated Movement Task

To generate high channel count datasets, we simulated offline datasets of a virtual user performing a target acquisition task (equivalent to the finger movement task). The goal of these simulations was to test the relative impact of then number of channels and amount of training data on decoder performance, rather than measuring absolute performance. The simulated user moved with a velocity proportional to the distance to the target along each DoF, with a random reaction time of 32-96 ms at trial onset. We generated artificial neural activity such that each channel had a random relationship with position, velocity, and acceleration, using the log-linear approximation suggested in ^51^:

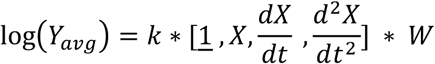

where *y_avg_* is a vector of average neural channel values *X*, is a vector of the current positions, *W* is a matrix of uniform random values between -1 to 1 and defines the random tuning of each channel, and (*W* is scaling constant adjusting the level of nonlinearity. At each time bin, the value of each channel was sampled from a normal distribution:

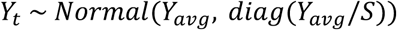

Where *y_t_* is a vector of values for each neural channel, and *S* is a constant that scales the noise standard deviation. Each channel had independent noise from other channels. We chose a value of *S* > = 10 such that the resulting LSTM decoding had similar accuracy to Monkey N (correlation of ∼0.8 at 100 channels). An additional lag of 32ms was added to the final positions and velocities relative to the artificial neural data. To account for random variation in simulated channel tuning, we ran multiple simulations for each analysis. We note that most simulated channels had non-zero tuning to kinematics, which may be unrealistic compared to real neural recordings where many channels may be untuned to the task.

Unless otherwise noted, simulated datasets used 1000 neural channels, 5 output DoFs, 10 min of data with 32 ms bins, and random target positions with a hold time of 500 ms.

### Speech Dataset

To test channel selection on a non-movement classification task, we used the speech decoding competition dataset from Willett et al. 2023^5^. One participant with Amyotrophic Lateral Sclerosis was implanted with four 64-channel Utah microelectrode arrays in cortical area 6v and area 44 (Broca’s area), for a total of 128 channels. Threshold crossings and spike band power (Nason 2020) features were extracted for each channel and binned in 20ms bins. Neural activity was recorded while the participant attempted to speak sentences, for a total of 12,100 sentences across 8 sessions. Here, we used neural features to decode phonemes and report the phoneme error rate. We note that a language model could be used (as in ^5^) to decode words and further improve error rates, however, here we focus on optimizing the decoder pre-language model for simplicity.

### Neural Network Decoders

We tested three neural network decoder architectures, chosen for their high offline or online decoding performance in other studies. Decoders took in binned neural features (spike events, threshold crossings, and/or spike band power depending on dataset) and were trained to predict the position and velocity of each degree of freedom. For the speech dataset, the decoder was trained to predict the most likely phoneme. The neural network decoders were implemented and trained using PyTorch 1.12.1.

### Recurrent Neural Networks

We tested two variants of recurrent neural networks: the long short-term memory (LSTM)^52^ which has previously demonstrated high decoding performance^35,53,54^, and a simple ‘vanilla’ recurrent neural network (termed ‘RNN’ here) which represents a simpler architecture with strong potential for compressibility. The LSTM was implemented using the *torch.nn.LSTM* class with a single fully connected (linear) output layer. The RNN was implemented using the *torch.nn.RNN* class using the ReLU nonlinearity, and used a single fully connected (linear) output layer. Each decoder took in the current time bin of neural features and the previous hidden state, updated the hidden state, and predicted outputs as a linear function of the hidden state. During training, a sequence length of 20 and zero-initialization for the hidden state were used for each sample. The final output of each sample sequence was used to compute the loss. During online decoding, the hidden state was stored in memory, such that the inputs at each timestep were the current neural bin and the previous hidden state.

### Temporal Convolutional Network (TCN)

We tested the TCN as described in Willsey et al. 2022^28^ and Temmar et al. 2024^55^, which has previously demonstrated strong real-time controllability^28,35^. The network uses an initial convolutional layer over time (for each channel) followed by several feedforward layers that use batch normalization, dropout, and ReLU. The network used 5 bins of time history per channel (160 ms). In contrast to the implementation in Willsey et al. 2022, here we normalized neural inputs, did not use the final batchnorm layer, and did not perform ReFIT recalibration for online trials.

### Decoder Training

For offline decoders, datasets were split into 70% for training, 10% for validation, and 20% for testing. For online decoders, datasets were split into 80% training (∼400 trials) and 20% validation (∼100 trials) since a test set was unnecessary. Neural features and output positions/velocities were normalized based on the training set to have zero mean and unit variance.

We first performed hyperparameter optimization for each decoder architecture on each dataset. We performed a random search over hyperparameter values using a minimum of 400 models trained per search. We chose the model size with minimal size that achieved maximal accuracy, and decoder performance was generally robust to the specific choice of hyperparameters. See Supplemental Table 2 and 3 for parameter values used. For the finger-brain and finger-EMG decoders, one additional day for each (separate from the days used for analysis) were used for hyperparameter search. For the reaching dataset, the train and validation sets were used for hyperparameter search. For the simulated datasets, additional datasets were generated for hyperparameter search.

Decoders were trained using minibatch gradient descent with the PyTorch Adam optimizer using the mean-squared error (MSE) loss. The learning rate was linearly decayed using a fixed iteration count. For online decoders, after training was complete, a linear regression was calculated to transform the normalized outputs back to the initial position and velocity scale. Training took less than 5 minutes for each movement decoding network.

### Speech Decoder Training

Speech decoder models used the same architecture and training method as that of Willett et al. 2023, but with modifications to perform channel selection. The core model used a 5-layer gated recurrent unit (GRU) with a linear output layer to predict the most likely phoneme at each timestep and was trained using a connectionist temporal classification (CTC) loss. A gating layer was added to the input (see below) to perform channel selection with a separate gate for each day, and an L1 penalty on the gate values were added to the connectionist temporal classification loss. The decoder implemented in PyTorch 1.12.1. The speech dataset was split into 70% training, 10% validation, and 20% testing. We performed a hyperparameter search over the learning rate, L1 penalty, L2 decay, and dropout, keeping all other hyperparameters at the defaults in Willett et al. 2023 (see final values in Supplemental Table 3). The hyperparameter search was performed with 80% of channels pruned, to better optimize for accuracy with few active channels.

### Pruned Decoder Training

Pruned decoders were trained by iteratively removing (zero-ing) parameters with the smallest magnitude values. No weights were pruned during an initial warmup period of 50 training iterations. Then, the fraction of pruned weights linearly increased from 0% to the final desired prune percentage. Parameters were then finetuned for a specific number of iterations without any additional pruning. The *torch.nn.utils.prune.l1_unstructured* module was used to prune parameters. Pruning was performed globally such that different layers could have different fractions of weights pruned. Through initial experiments we found that pruning the bias parameters and batchnorm layers (which represent a small fraction of total weights) often resulted in poor model performance or training instability, and thus did not prune those weights.

To achieve slightly better accuracy, we implemented pruning with rewinding as suggested in Frankle 2019. With rewinding, the decoder was first pruned normally to determine which weights remained unpruned after training (the “important” weights). The unpruned weights were then reset to their original random values (keeping the pruned weights at 0), and the decoder was retrained and optionally further pruned. Multiple steps of rewinding and pruning were repeated to reach the desired pruning level. We used pruning percentages of 50, 90, 95, 98, 99, 99.25, 99.5, 99.75. For example, if a final pruning percentage of 95% was desired, the model would be trained/pruned to 50%, rewound, to 90%, rewound, and to 95%. See Supplemental Figure 1 for example results comparing pruning methods.

### Sweeps over Model Sizes and Pruning Levels

To determine the optimal tradeoff between model compression and accuracy, we performed sweeps over model size and the pruning level. For the RNN and LSTM, the hidden size was varied from 32 to 512 units. The TCN hidden size (the same for every layer) was varied from 32 to 512 and the number of layers were varied from 1 to 5. For each size, the model was pruned between 0% and 99.7% percent. Correlation and MSE were computed for each size/pruning combination.

### Sparse Networks on an Embedded Microcontroller

To run neural decoders on implantable hardware without the need for customized hardware, we developed an embedded-C code library for efficiently running sparse networks. We tested with RNNs which could be implemented with few matrix operations and had the highest compressibility (see Results). As an overview, a single RNN update step was implemented as:

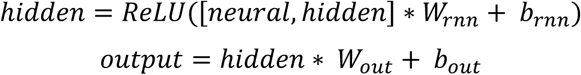

where *neural* contains the current neural features, *hidden* contains the hidden state, *W_rnn_/b_rnn_* are the RNN parameters, and *W_out_/b_out_* are the output layer parameters. *W_rnn_* and *W_out_* were > 90% sparse so that they could be efficiently stored and operated on using a sparse format. The pruned RNN decoder was trained normally in PyTorch and weights were exported to file, saved as value/matrix location pairs. Then, the microcontroller loaded the weights from an external SD card and a hidden state vector was initialized to zeros. To perform decoding, the microcontroller received binned neural features over a serial connection, performed a one timestep decoder update by using sparse matrix multiplication and addition, and outputted the kinematic value predictions. The hidden state was stored in memory for the next timestep. All computations were performed using 32-bit floating point values (although further speedup may be achieved by using quantizing to smaller values). The decoder latency was calculated as the total on-device processing time after receiving the current timestep of neural features and was measured to the nearest microsecond using the on-device clock. Latency was averaged over 100 decoding timesteps. We note that the latency measured here only includes the decoding processing time, and not time related to feature processing and binning.

We tested using the Teensy 4.1 microcontroller platform, which comprises a 32-bit 600MHz ARM Cortex-M7 processor with 8 Mb of native flash, 1024 Kb of RAM, and a floating-point math unit. Decoding latency was measured using a pruned RNN decoder with 96 neural inputs, a hidden size of 128, 95% pruned, and 1.7k parameters. Offline neural data was from the finger-brain dataset, with 96 neural channels and 4 kinematic outputs.

### Compression Factor Calculation

Model compression factors were calculated by dividing the number of parameters in the non-pruned (default) model by the number of non-zero parameters in the pruned model. For the compression factors in Figure 3, the default model was an LSTM with the hidden size chosen during hyperparameter search (where the LSTM achieved the overall highest accuracy for each dataset), and the pruned model was the RNN with the fewest non-zero parameters that maintained the desired accuracy level.

### Channel Selection

To perform channel selection with the neural network decoders, we added an input “gate” layer that aimed to learn the importance of each channel. Each input channel was multiplied by a single gate parameter, where values were enforced to be in the range 0 to 1 by applying a sigmoid function:
*gates* = torch.nn.Sigmoid()(*scalar* * *gates_pre*) where gates is the final value multiplied by each input channel, *gates_pre* is a learnable parameter, and *scalar* is a hyperparameter to adjust the sensitivity of the output values. *gates_pre* was initialized using a normal distribution with standard deviation 1e-6, for near-zero values with a small amount of randomness to break symmetries. Input values were multiplied by the *gates* and then passed through a batchnorm layer which was found to improve decoder accuracy. L1 regularization was applied to the *gates* to encourage channels to be masked to 0. Decoders were trained the same as the pruned decoder training, but only the gate layer was pruned. Thus, during training the gates learned the channel importance values and the least important channels were gradually pruned. Separate decoders were trained for each desired fraction of active channels, for maximum accuracy. All channel selection decoders for the movement datasets used the LSTM architecture. The loss function was the standard MSE loss with an added L1 penalty on the input gates:

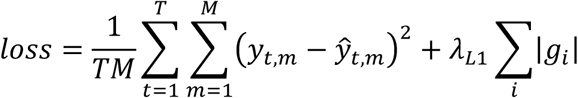

where *y* contains kinematic predictions with *T* timesteps and *M* outputs, *ŷ* contains the predicted kinematics, *λ_L_*_1_ is a hyperparameter to control the strength of the gate L1 penalty, and *g* is the weights of the input gates.

For the speech decoding models, a single gate parameter was used for both the threshold crossing and spike-band power features (both features from a channel were multiplied by the same value), the batchnorm layer after the gate was omitted (which resulted in higher accuracy), and independent gate parameters were learned for each day.

We evaluated pruning-based channel selection against random selection and linear channel selection. For random selection, a random subset of the gate values were set to zero before training and no additional channels were pruned. For linear selection, we performed a ridge regression between each channel and the kinematic outputs and calculated the average correlation. The channels with the lowest correlations were then removed by setting the decoder gate values to zero before training.

### Multi-Channel-Gate Decoder

To allow a single decoder to use varied numbers of active channels, we added multiple channel gates to the decoder (Figure 7a). For each desired number of active channels, a channel gate and linear layer was added to the input of the decoder. During decoding, features were passed through only one channel gate before the main decoder model. Channel gates were simultaneously trained with different final pruning levels.

### Online Decoder Testing

Sparse decoders were tested in online (real-time) trials with Monkey N on the finger movement task. Neural data was sampled and filtering using a Cerebus neural signal processor (see details above), sent to the xPC to bin and calculate features, which were then sent to a computer for decoding which predicted finger position(s). These positions for each finger were sent back to the xPC to update the virtual hand (see ^28^ for more details). Decoders were run in Python 3.9 using a dedicated Linux computer with an RTX 2070 Super GPU (NVIDIA). The final displayed position was a combination of predicted velocity and position (98% integrated velocity, 2% position) which helped prevent the virtual fingers from becoming biased toward one direction (similar to that of ^56,57^)

During online comparisons, decoders were alternated in an A-B-A-B format with a minimum of 100 trials per set, with performance averaged across sets. An online decoder test was stopped if the monkey was unable to acquire targets for more than 30 seconds or if the monkey stopped attempting to acquire targets. Online performance was measured using trial time which includes both the movement and hold periods. The first 50 trials of each set were omitted from the performance calculation to account for the user’s learning of the decoder. On two days the monkey was unable to acquire targets near the boundaries of the range (far extension or far flexion), and these trials were excluded for all decoders on the day (see Supplemental Table 1 for day specific information).

To test generalization of sparse decoders to unseen target positions, we trained the decoders on open-loop trials with only 9 target positions (extend-rest-flex for two finger groups). The trained decoder was then tested in real-time on the full range of target positions (randomly chosen throughout the full range).

## Supplemental Pruning Results

**Supplemental Figure 1:**
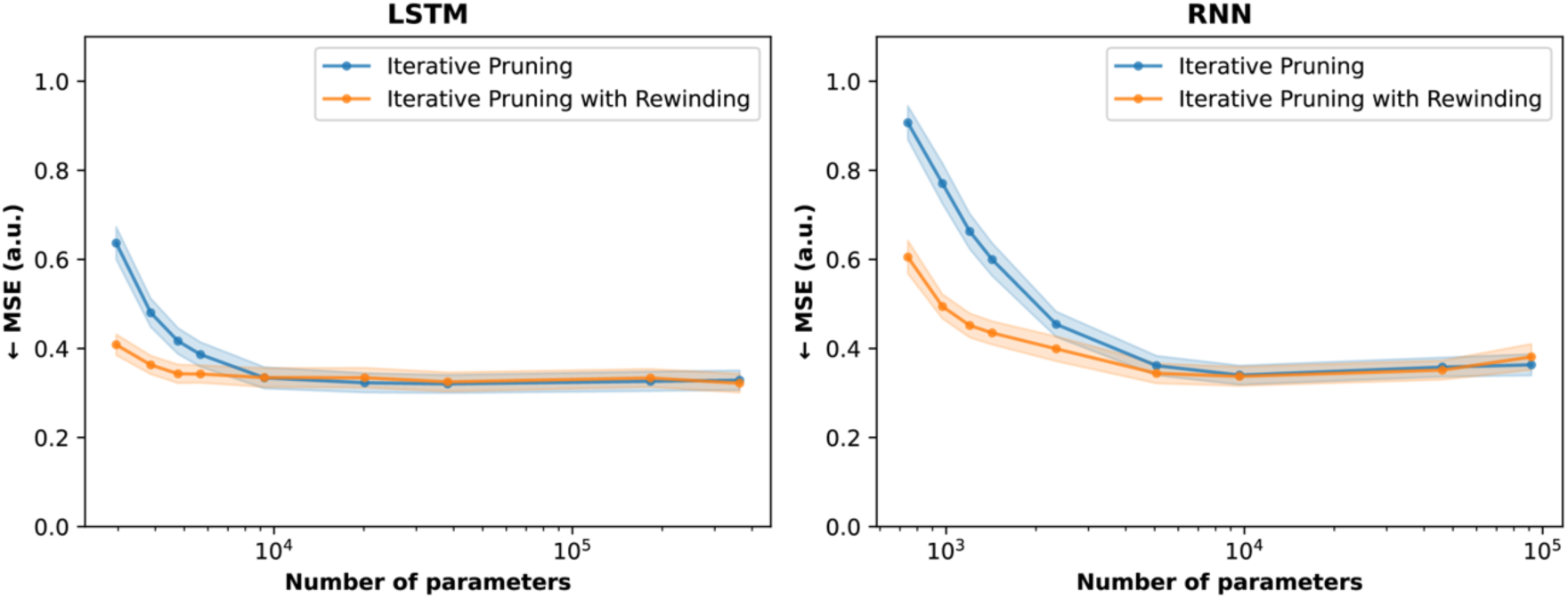
Offline accuracy for iterative pruning with and without rewinding. Iterative pruning with rewinding achieves better accuracy (lower MSE) at low parameter counts, allowing for slightly higher decoder compression. Hidden size was held constant at 256 and the fraction of pruned weights was varied from 0% to 99.75%. Shaded error bars denote 1 S.E.M, averaged over the 10 intracortical finger datasets. Iterative pruning gradually increases the fraction of weights pruned during training, ending with the desired pruning fraction. Iterative pruning with rewinding (as shown in Figure 2 and described in Methods), gradually prunes to given fraction, resets the non-zero weights to their original values, and continues training and pruning to a higher fraction of pruned weights. This rewinding process can be performed multiple times, which first allows the network to learn which weights will remain non-zero and then allows the weights to be finetuned.

**Supplemental Figure 2:**
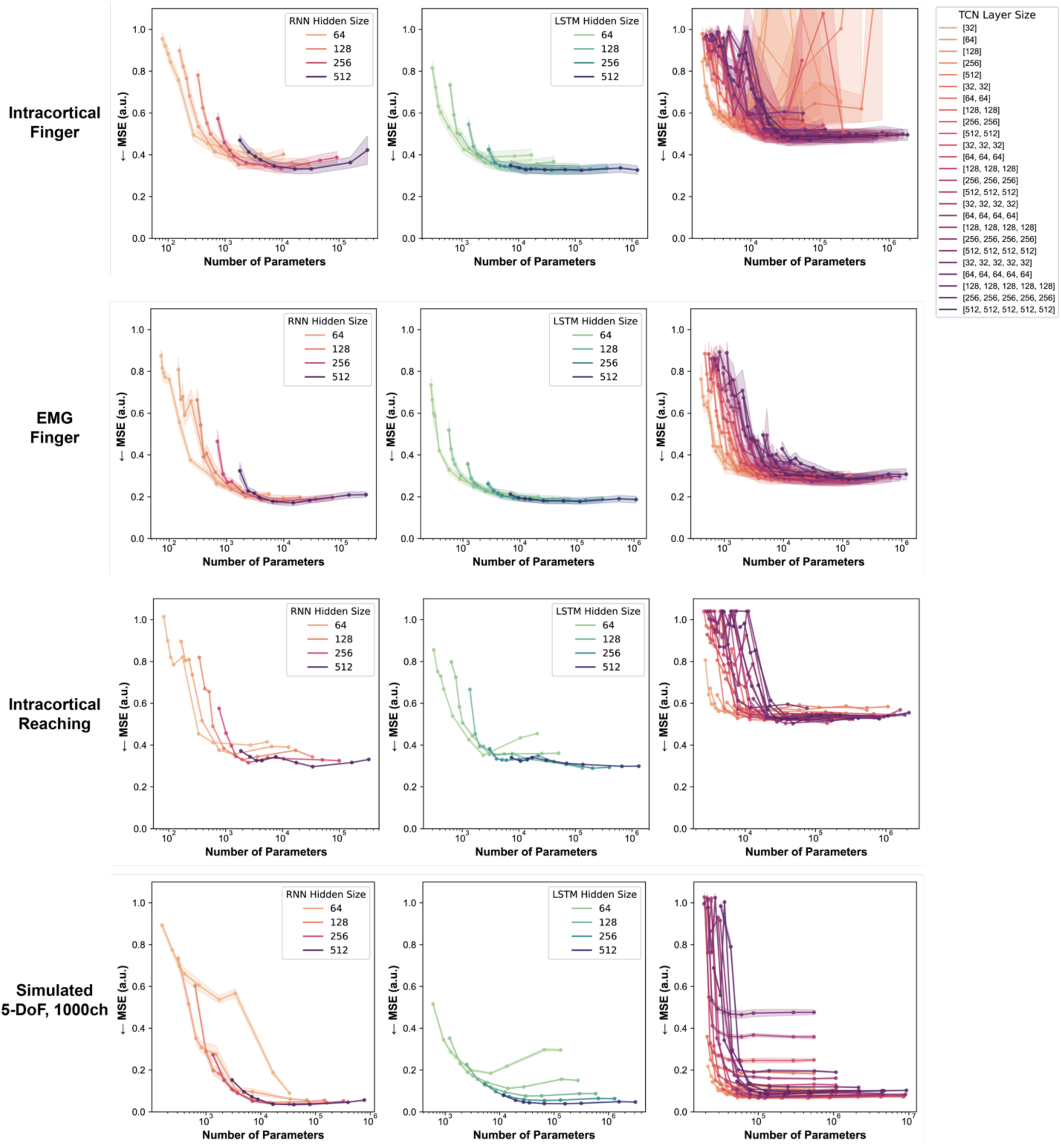
Offline accuracy for all models across varied hidden sizes and varied fraction of pruned weights. Going from right to left along each line moves from 0% to 99.75% pruning. Left – RNN, Middle – LSTM, Right – TCN. For the RNN (left column), pruning by a moderate amount results in slightly improved MSE compared to no pruning (the no-pruning case can be found be the right-most point on each colored line). The curves in Figure 3e-h (Pareto boundary curves for accuracy vs number-of-parameters) were calculated by finding the lowest MSE for each number-of- parameters in the data shown here. Shaded error bars denote 1 S.E.M, averaged across datasets. The TCN layer size legend lists the size of each layer in the network, where “[256, 256, 256]” indicates a three layers each with size 256.

**Supplemental Table 1:**
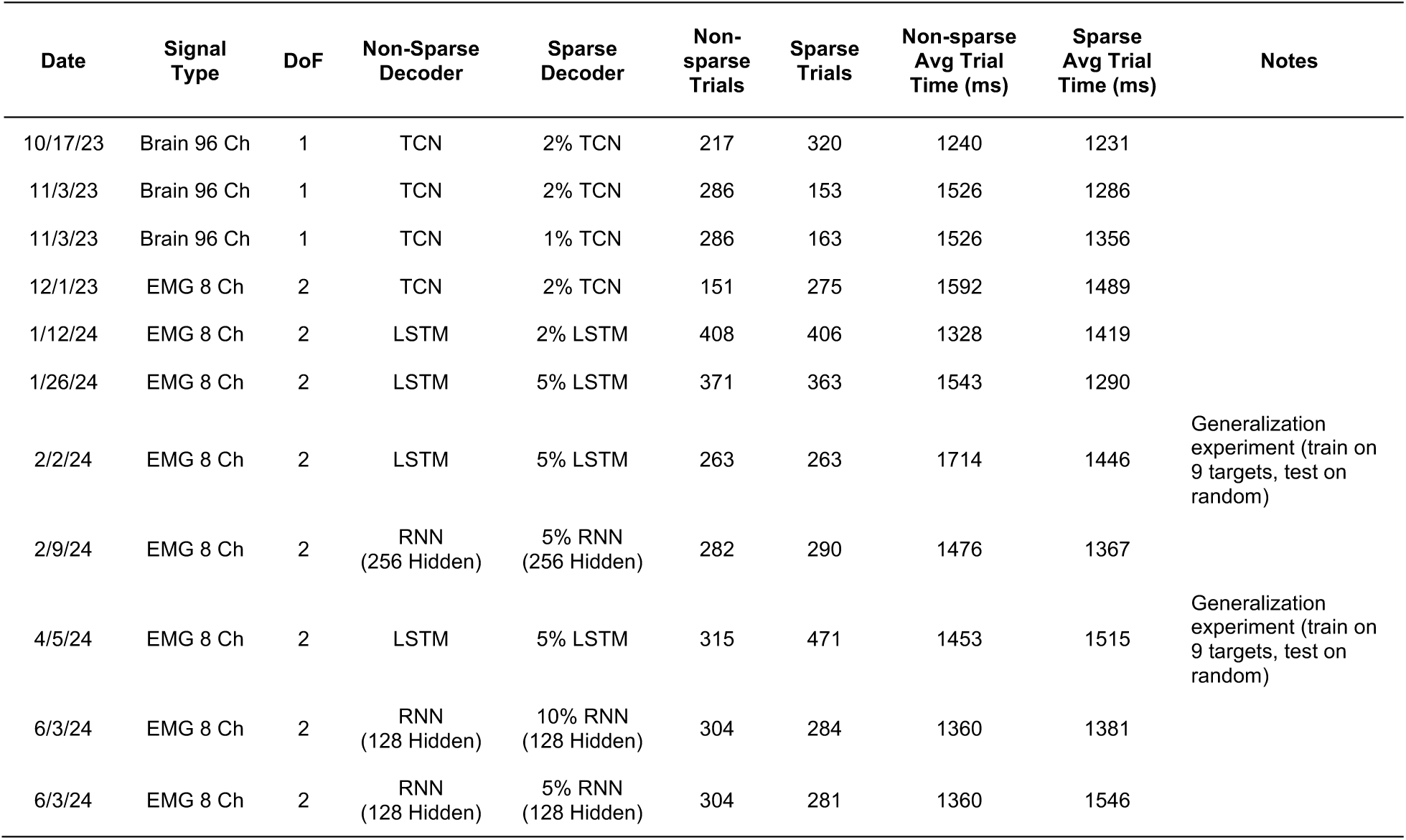
Online experiment sessions using sparse decoders. Sessions were with Monkey N and consisted of approximately 10 minutes of open-loop (offline) data collection, training a normal and one or more sparse decoders, and testing the decoders in closed-loop (online) trials.

| Date | Signal Type | DoF | Non-Sparse Decoder | Sparse Decoder | Non-sparse Trials | Sparse Trials | Non-sparse Avg Trial Time (ms) | Sparse Avg Trial Time (ms) | Notes |
| --- | --- | --- | --- | --- | --- | --- | --- | --- | --- |
| 10/17/23 | Brain 96 Ch | 1 | TCN | 2% TCN | 217 | 320 | 1240 | 1231 |  |
| 11/3/23 | Brain 96 Ch | 1 | TCN | 2% TCN | 286 | 153 | 1526 | 1286 |  |
| 11/3/23 | Brain 96 Ch | 1 | TCN | 1% TCN | 286 | 163 | 1526 | 1356 |  |
| 12/1/23 | EMG 8 Ch | 2 | TCN | 2% TCN | 151 | 275 | 1592 | 1489 |  |
| 1/12/24 | EMG 8 Ch | 2 | LSTM | 2% LSTM | 408 | 406 | 1328 | 1419 |  |
| 1/26/24 | EMG 8 Ch | 2 | LSTM | 5% LSTM | 371 | 363 | 1543 | 1290 |  |
| 2/2/24 | EMG 8 Ch | 2 | LSTM | 5% LSTM | 263 | 263 | 1714 | 1446 | Generalization experiment (train on 9 targets, test on random) |
| 2/9/24 | EMG 8 Ch | 2 | RNN (256 Hidden) | 5% RNN (256 Hidden) | 282 | 290 | 1476 | 1367 |  |
| 4/5/24 | EMG 8 Ch | 2 | LSTM | 5% LSTM | 315 | 471 | 1453 | 1515 | Generalization experiment (train on 9 targets, test on random) |
| 6/3/24 | EMG 8 Ch | 2 | RNN (128 Hidden) | 10% RNN (128 Hidden) | 304 | 284 | 1360 | 1381 |  |
| 6/3/24 | EMG 8 Ch | 2 | RNN (128 Hidden) | 5% RNN (128 Hidden) | 304 | 281 | 1360 | 1546 |  |

## Decoder Hyperparameters

**Supplemental Table 2:** Hyperparameters for sparse decoders. Values were optimized using a validation set as described in Methods. Note that the recurrent models processed the sequence length (“history bins”) in a sequential manner, whereas the TCN used the entire sequence at once. For the TCN, the “Num Layers” is the number of feedforward layers in addition to the convolutional input layer.

|  | Intracortical Finger |  |  | EMG Finger |  |  | Intracortical Reaching |  |  | Simulated 5-Dof, 1000ch |  |  |
| --- | --- | --- | --- | --- | --- | --- | --- | --- | --- | --- | --- | --- |
|  | LSTM | RNN | TCN | LSTM | RNN | TCN | LSTM | RNN | TCN | LSTM | RNN | TCN |
| History Bins | 20 | 20 | 5 | 20 | 20 | 5 | 20 | 20 | 5 | 20 | 20 | 5 |
| Num Layers | 1 | 1 | 4 | 1 | 1 | 4 | 1 | 1 | 3 | 1 | 1 | 3 |
| Hidden Size | 256 | 256 | 256 | 200 | 150 | 256 | 256 | 256 | 256 | 512 | 256 | 1200 |
| Start Learning Rate | 1.00E-03 | 5.00E-04 | 1.00E-03 | 1.00E-03 | 5.00E-04 | 5.00E-04 | 1.00E-03 | 5.00E-04 | 2.00E-04 | 5.00E-04 | 5.00E-04 | 2.00E-04 |
| End Learning Rate | 1.00E-05 | 1.00E-05 | 1.00E-04 | 1.00E-05 | 1.00E-05 | 1.00E-04 | 1.00E-05 | 1.00E-05 | 1.00E-04 | 1.00E-05 | 1.00E-05 | 1.00E-04 |
| Weight Decay | 1.00E-04 | 1.00E-04 | 2.00E-03 | 1.00E-04 | 1.00E-04 | 1.00E-04 | 1.00E-04 | 1.00E-04 | 1.00E-03 | 1.00E-04 | 1.00E-04 | 1.00E-04 |
| Dropout | - | - | 0.50 | - | - | 0.05 | - | - | 0.40 | - | - | 0.10 |
| Iterations | 2000 | 2000 | 2000 | 2500 | 2500 | 3000 | 2500 | 2500 | 2500 | 2500 | 3000 | 3000 |
| Batch Size | 64 | 64 | 64 | 64 | 64 | 64 | 64 | 64 | 64 | 64 | 64 | 64 |

**Supplemental Table 3:** Hyperparameters for the channel selection decoders. Values were optimized using a validation set as described in Methods. For the speech classification decoder all hyperparameters (including those not listed here) were the same as in Willett et al. 2023^1^, except for learning rates, L2 weight decay, dropout, and the gate L1 weight, which were optimized using a validation set.

|  | Intracortical Finger | Intracortical Reaching | Simulated 5-Dof, 1000ch | Speech Classification |
| --- | --- | --- | --- | --- |
|  | LSTM | LSTM | LSTM | GRU |
| Num Layers | 1 | 1 | 1 | 5 |
| Hidden Size | 400 | 300 | 1200 | 1024 |
| Start Learning Rate | 2.00E-04 | 2.00E-04 | 2.00E-04 | 2.00E-02 |
| End Learning Rate | 1.00E-06 | 5.00E-06 | 1.00E-05 | 1.00E-02 |
| Weight Decay | 5.00E-04 | 2.00E-04 | 1.00E-04 | 1.00E-05 |
| Iterations | 3000 | 1500 | 3000 | 10000 |
| Gate L1 Weight | 0.2 | 0.2 | 0.2 | 1.00E-05 |
| Batch Size | 64 | 64 | 64 | 64 |
| Dropout | - | - | - | 0.20 |

## Considerations for Decoding On- vs Off-Implant

Wireless, implantable BMIs can either (1) transmit neural features to an external computer to perform decoding or (2) perform sparse decoding on the implant and transmit decoded control signals to external devices (e.g., robotic prostheses, computers, phones). Here we evaluate multiple factors for choosing between the two options. We consider a 1000-channel system and a sparse decoder with 7k parameters.

### Power Consumption

For option (1), we assume the system will transmit low-data-rate features, for example, 20 ms (50 Hz) binned features at 8-bit resolution, or 400 bps per channel. Assuming a worst-case overhead of 100% (such as for packet headers and error correction), this leads to 800 bps per channel, or 800 kbps per 1000 channels. Using 8.5 pJ per transmitted bit^2,3^ this results in an estimated 6.8 µW of power.

For option (2) we would like to estimate the optimal power consumption for running a sparse decoder on an embedded system. However, power consumption can significantly vary based on system architecture and the availability of custom hardware accelerators. As a lower bound on power consumption, the optimized decoding system of An & Nason-Tomaszewski et al. 2022^4^ used 45 or 17 µW (for an M0 core or a hardware accelerator, respectively) for a decoder with 192 parameters. As an upper bound, the system of Liao et al. 2024^5^ decoded using 600 µW for a neural network with 526k parameters. Thus, the sparse decoder proposed here using 7k parameters could likely be run using much less than 500 µW, and likely less than 50 µW (when using optimized accelerators with >1T operations/W ^6^). While this is significantly greater than option (1), it still may be within an implant’s power budget, where 50 µW would only increase the power consumption of low-power, high-channel count recording chips (such as the 24.7mW chip in Yoon et al. 2021^7^) by less than 0.3%.

### Data Availability and Privacy

Option (1) allows for easy access to neural features by an external computer. For collecting training data to improve decoders (both across users and within users), this may be necessary. However, if this data is not necessary (for example, if decoder recalibration is not needed or recalibration can be performed on-implant), then option (2) provides enhanced data privacy and security for the user, both in terms of exposing their neural activity to the external computer and the potential for eavesdropping on data over a wireless link. One could also imagine using a “training” mode in which data is transmitted off-device in order to train a decoder, and then switch running the decoder on-device, option (2), for enhanced data privacy.

### Decoding Latency

Previous work has found that lower-latency decoder updates and feedback can improve closed-loop BMI control^8–10^. Wireless data transmission may have increased delays relative to wired connections. If the BMI’s output device is a computer or phone that requires receiving wireless data or commands, then there may only be a small latency difference between options 1 – transmit neural data (∼1.6 Mbps) and decode on the computer, and 2 – decode on implant and transmit commands (both options only require 1 transmission). However, for controlling other devices such as a robotic prosthesis, option (1) may require transmitting data to a computer for decoding followed by a second transmission to the prosthesis, whereas option (2) would require a single transmission of commands directly from the implant to the prosthesis. Here, option (2) only requires a single transmission and may have lower latency. Future BMIs may also provide sensory stimulation for feedback; if stimulation is in-part driven by the decoder, then the lower latency of option (2) may provide enhanced feedback by eliminating round-trip wireless transmission to an external computer.

### Decoder Updates

The decoder may need to be updated or retrained to account for shifts in the neural activity over time. For option (1), this may be easier since recent neural data is readily available on the external computer and the computer may have more processing power and memory to process updates or loading multiple decoders. For option (2), this would either require strategies for updating weights on-device (for example, updating channel-wise normalization prior to decoding may be simple) or receiving new weights from an external computer.

### Summary

While option (1) has slightly lower power consumption, the power consumption for data transmission or sparse decoding has minimal impact on the total power consumption of the implantable BMI, making either option viable. For ease of decoder updates, option (1), decoding externally, may be advantageous. For improved data privacy and lower latency, option (2), decoding on-implant, may be better.

## Estimating Power Savings from Deactivating Channels

Here we aim to estimate how much power could be saved on implantable BMIs by deactivating uninformative recording channels. In the main-text we found that 50-89% of channels could be deactivated with minimal loss in decoding accuracy. The fraction of channels that can be deactivated likely depends on the signal quality and degree of tuning, the total number of channels, and the overall task complexity.

For this estimation, we assume that amplifiers for individual channels can be deactivated, eliminating their power consumption. We ignore any power savings from the analog-to-digital converter (ADC) which is often multiplexed between multiple channels (although with fewer channels, a lower clock speed could likely be used for slightly lower power).

In low-power BMI systems, amplifiers typically consume 87-94% of the total system power (^3,4,11^; ignoring the optical-related power in ^11^). For this estimation we use 90%.

With 50% of channels removed, the total power is 55% of the original, for a 1.8x power reduction. With 89% of channels removed, the total power is 20% of the original, for a 5x power reduction. Thus, channel deactivation could result in approximately 1.8-5x power savings. While this specific number will vary based on the BMI-related factors mentioned above and the specific hardware implementation, the technique of channel-deactivation could be used across a variety of recording hardware designs.

## Few-Channel Decoder Robustness to Noise

When deactivating large proportions of channels, one might expect the model to rely more on each remaining channel and be less robust to noise or other channel perturbations. Here we evaluated few-channel decoder offline accuracy for different types and amounts of noise. We tested the following types of noise:

- ***Bias Noise:*** A random bias is added to each channel, constant across time and chosen from a Gaussian distribution. This simulates systematic offsets in channel readings, such as baseline activation drift.
- ***Scaling Noise:*** A multiplicative noise is applied to each channel, with scaling factors sampled from a Gaussian distribution. This type of noise models scenarios where channel gain fluctuates.
- ***White Noise:*** Independent Gaussian noise is added to each time point of each channel. This simulates general noise sources such as thermal noise or increased neural spiking variability.
- ***Random Walk Noise*:** A random walk is added to each channel by summing Gaussian noise across time, resulting in cumulative, drifting deviations. This simulates noise processes like slow electrode drift.
- ***Channel Dropout:*** Entire channels are randomly zeroed out (with dropout probability controlled by the noise parameter) and remain constant across time. This mimics transient or persistent channel failures, such as those caused by electrode disconnection or hardware malfunction.

For this analysis we used the neural-finger dataset with 96 channels. We trained a baseline LSTM (using all channels) and a few-channels LSTM (using 11% of channels) on each day (10 days total), and evaluated offline correlations using a test set when noise was added to the neural inputs.

Noise was added after neural data was binned. For the channel-dropout noise, the dropout probability was across all channels, not just the active channels. Note that each neural channel was normalized to have unit standard deviation (based on a separate training set) prior to adding noise.

As seen in Supplemental Figure 3, across all types of noise, both models have a similar drop in accuracy as the noise level is increased. If one model was less robust to noise, we would expect a larger accuracy drop compared to the other model. For scaling-noise and for channel-dropout there was a slightly larger drop in accuracy from the few-channel decoder, as seen by the larger gap between lines. However, for each noise level and each type of noise, no significant differences in average correlation were found between the two decoders (two-sided t-test with Bonferroni correction). Thus, reducing the number of active channels does significantly reduce the robustness to noise (at least in the open-loop condition).

**Supplemental Figure 3:**
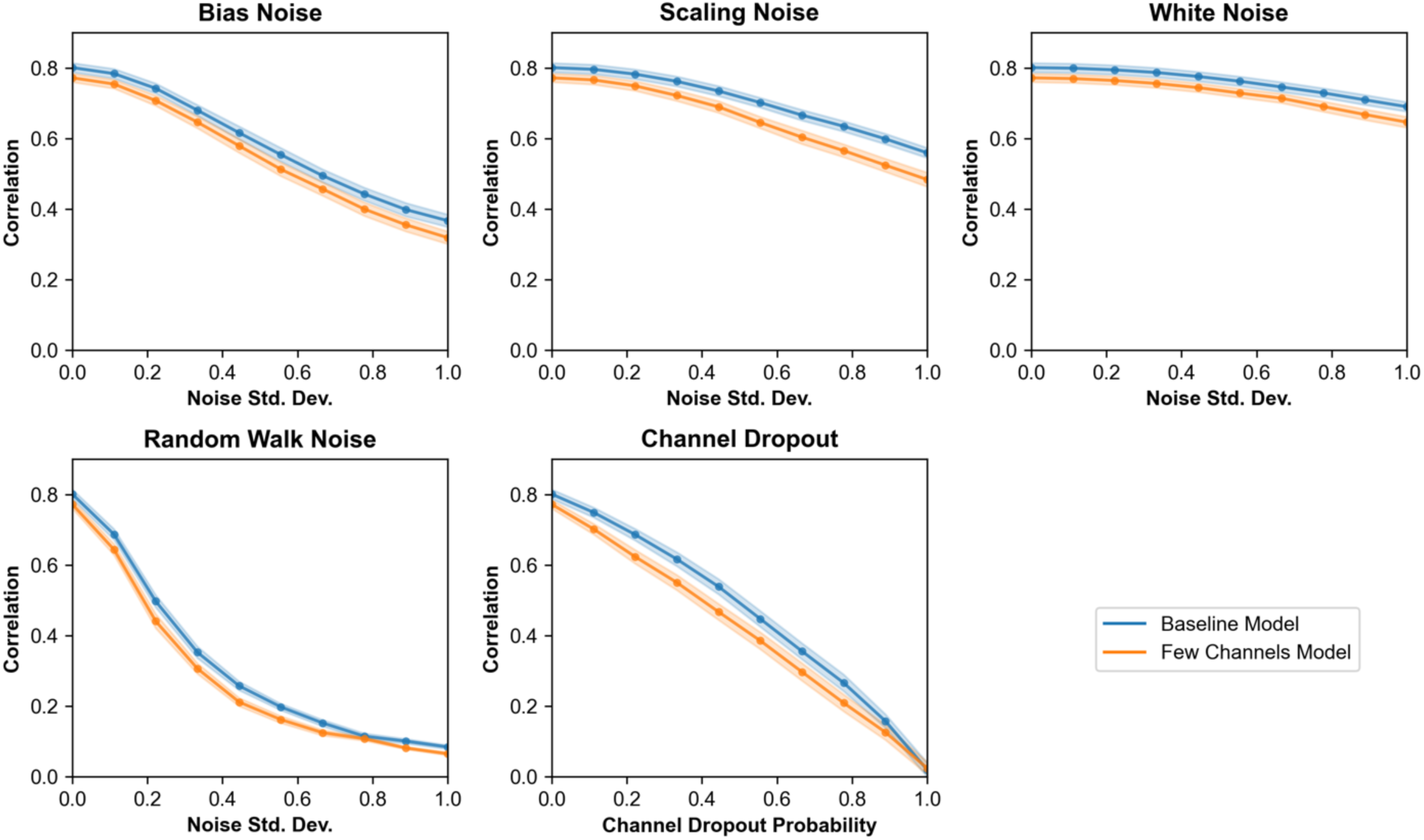
Few-channel decoder robustness to neural noise. The baseline model used all available channels and the few-channels model used 11% of total channels. Results are for the neural finger task (96 total channels) and averaged across 10 datasets, where a new set of models were trained for each dataset. Correlation is between the true and predicted kinematics (velocity and position) using a held-out test set. Error bars represent 1 S.E.M. No significant differences in correlation were found between the two decoders across all types and magnitudes of noise (two-sided t-test with Bonferroni correction).

